# Assignment of frost tolerant coast redwood trees of unknown origin to populations within their natural range using nuclear and chloroplast microsatellite genetic markers

**DOI:** 10.1101/732834

**Authors:** Natalie Breidenbach, Oliver Gailing, Konstantin V. Krutovsky

**Author notes:** Corresponding author (K.V.K).

## Abstract

Considering climate change and expected changes in temperature and precipitation, some introduced timber species are prospective for growing in Germany or Europe to produce valuable wood products and support sustainable forestry. The Californian tree, coast redwood (*Sequoia sempervirens* [D. Don] Endl.) is one of such species due to its excellent wood properties and high growth rate. It is sensitive to the freezing temperatures, but several trees of unknown origin introduced to Germany decades ago demonstrated high frost tolerance, and their propagated cuttings were planted all over German botanic gardens and arboreta. The knowledge of their origin within the natural distribution range could help us identify the potential genetic resources of frost resistant coast redwood genotypes. Therefore, both trees of unknown origin in Germany (G) and two reference data sets representing the “Kuser provenance test” established in 1990 in France (F) and samples collected in California (C) with known origin were genotyped using 18 microsatellite markers including 12 nuclear and six chloroplast simple sequence repeat (cpSSR) markers. The number of haplotypes found in the data sets based on six cpSSR markers was surprisingly very high. These markers were used to assign the German frost resistant trees (G) to the two reference data sets (F and C). The genetic structure among California samples (C) based on nSSR and cpSSR markers was very weak and mainly due to northern and southern clusters separated by the San Francisco Bay as a geographic barrier between coast redwood populations confirming previously published data. It was impossible to confidently assign frost tolerant trees (G) to single native populations, but rather to either the northern or southern cluster. However, the existing frost tolerant genotypes can already be used to establish commercial coast redwood plantation for future German forestry.

## Introduction

Coast redwood (*Sequoia sempervirens* [D. Don] Endl.) is one of the four redwood species in the world. All of them are characterized by a particular red coloured wood, an endemic narrow natural distribution range and listed as threatened or endangered by the IUCN [1, 2]. Coast redwood stands out from the other three redwood species due to its valuable wood, high growth rate and natural vegetative reproduction [3–5]. It is a very important timber species in the natural range along the pacific coast in northern California and in the outside regions with similar climatic conditions established for timber production in several countries, such as France [6], New Zealand [7], and China [8]. The natural distribution range includes three plant hardiness zones 9b, 10a, and 10b (from −4 °C to 5 °C) categorized as humid Mediterranean climate [9]. Therefore, coast redwood usually experiences light frost only few weeks during the year [2] and is frost sensitive, especially at young age [3]. The coast redwood needles are capable to uptake water directly from the atmosphere [10], and coast redwood highly depends on frequent fogs that compensate the low precipitation during summer months [3, 11].

Increased precipitation and temperature in winter are expected in central Europe [12] including some regions in Germany and making its environment more suitable for some foreign tree species, such as coast redwood. Meanwhile, in some regions climate change will negatively affect native species [13]. Considering climate change, and the forecast for future climate, Germany forestry needs new adaptive management strategies [14]. To maintain German forestry and the growing demand for wood products in Europe more non-native tree species should be tested for potential introduction. One of the successful examples is the North American species Douglas-fir widely established in European forestry [15, 16]. The predicted altitudinal shift in the distribution of woody species [12, 17] due to climate change would make Germany more suitable also for species adapted to the Mediterranean climate, such as coast redwood.

Coast redwood has already been introduced to Europe as an exotic species [18]. Plant hardiness zones in Germany range from 7a (−17 to −15 °C) to 9a (−6 to −4 °C) [19] and present currently high risk for planting coast redwood as a timber species in Germany. The hardiness zones 8b (−12 to −9 °C) and 8a (−9 to −6 °C) along the Rhine valley in western Germany have climate conditions most similar to the natural distribution range of coast redwood, therefore most coast redwood trees were planted there. To mitigate climate change associated with predicted possible extreme droughts and low temperature occurrences in central Europe, German forestry needs to implement adaptive management based on frost and drought resistant genotypes of major timber species including introduced species such as coast redwood [6]. Information on their origins helps to select such genotypes. For instance, some of the coast redwood trees planted in the Rhine valley were frost resistant and survived temperatures as low as −20 °C [18], but their origins are unknown. The knowledge of the climatic conditions at their origins within the natural range could support this observation. Ideally, frost resistance should be also verified in controlled frost experiments.

Coast redwood is a hexaploid species and among only a few natural polyploids in gymnosperms [20–22]. It is a unique and valuable species, but its genetic structure is still insufficiently studied [23–30]. The earlier published results based on a range wide sample genotyped using six nuclear [31] and one chloroplast [30] microsatellite markers suggested high genetic diversity and low population differentiation with a weak indication of two clusters. It was demonstrated earlier that chloroplasts are strictly paternally inherited in coast redwood [32]. Coast redwood is prone to vegetative clonal propagation, and clone identification was possible for coast redwood with high confidence using the available microsatellite markers [29, 33]. Here we used available microsatellite or nuclear simple sequence repeat (nSSR) [26,31,33] and newly developed chloroplast SSR (cpSSR) and expressed sequence tag-SSR (EST-SSR) [34] markers to increase the resolution of population differentiation within the natural distribution range intending also to discriminate populations growing under suboptimal conditions.

The main objectives of this study were to 1) compare genetic population structure resolution of nuclear and chloroplast microsatellite markers, 2) study chloroplast microsatellite haplotype diversity based on a range-wide dataset, and 3) assign German frost resistant individuals to reference populations in California using cpSSRs.

## Material and methods

### Sample collection

Three data sets were used in this study: French (F), Californian (C), and German (G). The French data set F consisted of samples representing 153 trees from one of the international provenance test coast redwood planting site established in 1990 in St. Fargeau, central France by Kuser et al. [34] (so-called “Kuser provenance test”). It was established using seedlings representing 90 provenances of the natural distribution range of *S. sempervirens*. The individuals were propagated by cuttings and used to establish hedge orchards in California, Oregon, Spain, Britain, France, and New Zealand. In 2014, needle or cambium material of 153 surviving trees was collected in St. Fargeau and frozen until DNA extraction. In 1992, after a late frost occurrence in 1991, good frost tolerance was observed by the owner for 12 of the 158 collected trees (unpublished data). The data set F was compared with another “Kuser provenance test” experiment established in 1984 at a site, close to Berkeley, California, named the Russell Reserve. Douhovnikoff and Dodd [31] partitioned the Russell Reserve trees into 17 watersheds according to their original latitude location and the GPS coordination data (**Error! Reference source not found.**) for population genetic structure analysis based on the six nSSR markers [31]. To compare our data with their results and population based analyses [31], the St. Fargeau samples were also partitioned into 17 watersheds according to the mean latitude values for each watershed published in Douhovnikoff and Dodd [31] (S1 Table). Original locations of the ‘Kuser’ samples placed on the map with the mean monthly temperature pattern for the time period 1979-2013 are presented in S1 Fig.

**Fig 1.**
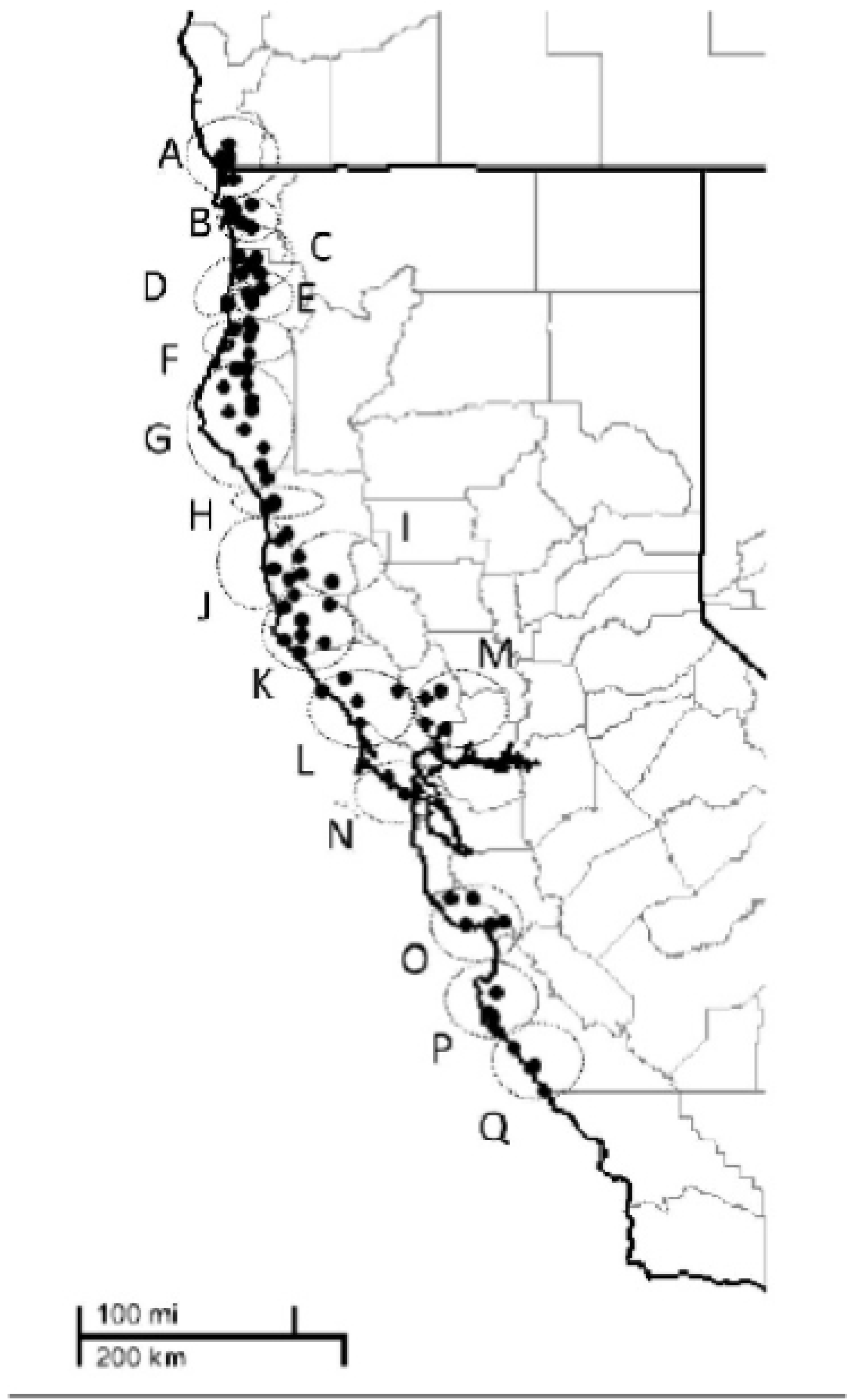
Map of 17 watersheds (A-Q) along the latitude within the natural distribution range of *Sequoia sempervirens* for the French data set F. (Fig 1 in Douhovnikoff & Dodd [31] reproduced with permission from The American Midland Naturalist).

The Californian data set C consisted of samples from 16 locations within the natural distribution range in central and northern California collected during the field trip in 2017. The locations were chosen according to their geographic position and the climatic conditions. Further, the original locations of the 12 frost resistant trees in the St. Fargeau provenance test site were considered in selecting potential sources for the frost resistant genotypes.

#### Acknowledgements

We thank the Bureau of Land Management of the Headwater Forest Reserve, Lynn Webb from the Department of Forestry and Fire Protection, Jackson Demonstration Forest, the Napa Land Trust, Napa Open Space District, Peter Leuzinger from Calfire (Sonoma-Lake-Napa Unit), the Pacific Union College, Enchanted Hills, Mt. Madonna County Park, East Bay Regional Parks District, and Candi Wozniak from the Redwood Ridge Estate for their permission to collect samples and their great support. We would like to thank Paul Asmuth, Bill Libby and Zane Moore for their suggestions and support during the sample collection in California. We thank Christof Niehues and his Baumschule Allerweltsgrün (Köln) for providing the samples from St. Fargeau and the clones for the genotyping validation. We thank Alexandra Dolynska, Christine Radler, Melanie Eckholdt and Babalola Jumoke for their help during lab work and Vadim Sharov (Genome Research and Education Center, Siberian Federal University) for help with computer analysis.

Unfortunately, one of the populations in southern Oregon was selected for sampling, but could not be visited during the field trip in 2017 due to forest fires. The optimal climatic conditions for *S. sempervirens* are along the coast, with high oceanic influence and humidity and high ground water level [35]. The selected locations were at higher altitude, more interior, and less foggy, and therefore represented drier and more extreme habitats [11, 36]. Sampled trees had a diameter at breast height (DBH) of at least 15 cm. The collected needle material was dried on silica gel. Number of samples, historic precipitation, and mean temperature per location are listed in S2 Table.

**Fig 2.**
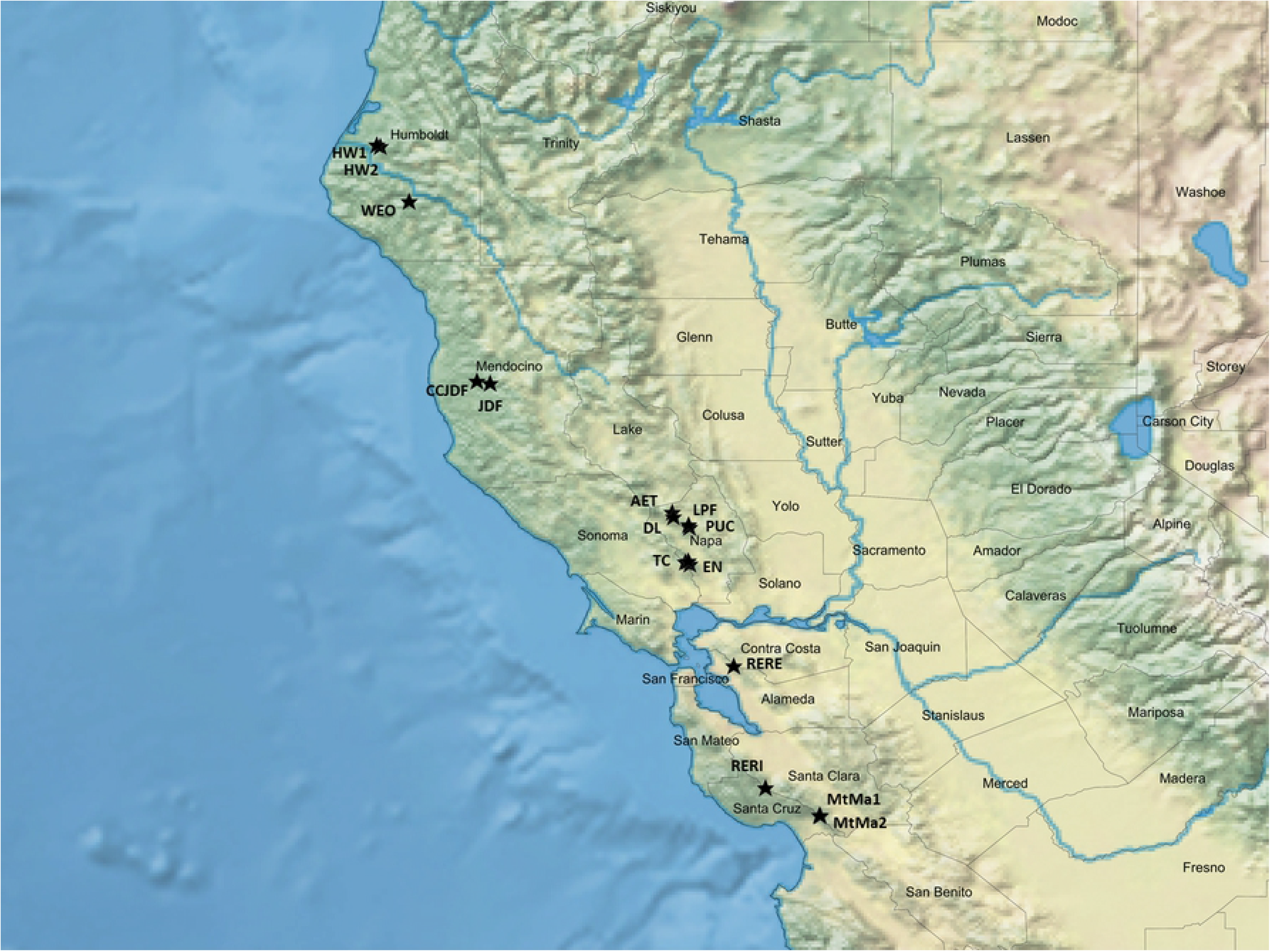
Collection sites for the Californian data set C depicted by asterisks in the California map generated using the SimpleMappr online software. (https://www.simplemappr.net).

The set was partitioned either into the 17 watersheds according to Douhovnikoff and Dodd [31] or two groups of northern and southern (south of the Napa Valley) watersheds, respectively.

The German data set G represented three collection sites: “Sequoiafarm Kaldenkirchen”, “Aboretum Burgholz”, and “Weltwald in Bad Grund” (S2 Fig). The “Sequoiafarm Kaldenkirchen” was established by the Martin family in the 1950’s using *S. sempervirens* seedlings representing likely a single tree from the Californian “Schenck-Grove”, Prairie Creek Redwoods State Park (Humboldt County, California, USA). Cuttings from these “best performing” clones were planted all over Germany and are known as the “Martin-Clones” (M. Geller, pers. communication). They were planted also at the two other sites, “Aboretum Burgholz” and “Weltwald Bad Grund”, in unknown numbers and combinations. The “Sequoiafarm Kaldenkirchen” and “Aboretum Burgholz” are both located in the Rhine-valley in the plant hardiness zones 8a and 8b [19], respectively. The “Weltwald Bad Grund” is an arboretum located in the western Harz Mountains near Bad Grund with relatively cold winters and dry summers, typical for the plant hardiness zone 7b. Additional individual trees were collected from botanic gardens in the following colder regions of Germany: Chemnitz (CH1), Bayreuth (BA030549 and BA 10067), and Göttingen (GOE_B, GOE_G, GOE_K, and GOE_P). Chemnitz and Bayreuth belong to the plant hardiness zone 7a, and Göttingen to zone 7b. Fresh needle material was collected from all these individual trees in Germany and frozen at −20 °C until DNA extraction.

Needles from 30 approximately three years old clones, including 2-3 ramets per clone, respectively, were collected in the Allerweltsgrün nusery (Köln) and frozen at −20 °C (S3 Fig).

### DNA isolation and PCR amplification of the microsatellite (SSR) markers

DNA was isolated from needles using the DNeasy Plant Kit (Qiagen, Hilden, Germany) following the manufacturer’s instructions. The isolated DNA was diluted in ddH_2_O 1:10 for PCR amplification and stored at −20 °C.

Twelve nuclear (five nSSR and seven EST-SSR) and six chloroplast (cpSSR) microsatellite markers were used in this study to genotype samples in all three data sets F, C and G (S3 Table). The same touch-down PCR program was used for all 18 PCR primer pairs following the protocol described in Breidenbach et al. [37]. The PCR products were separated and visualized using the ABI genetic analyser 3130xl with GENSCAN ROX 500 as an internal size standard.

### Verification of the nuclear microsatellite (nSSR) markers

The PCR primer nucleotide sequences for the nSSR markers were mapped to the *S. sempervirens* draft nuclear and chloroplast genome assemblies that become very recently publicly available to verify annealing sites for the microsatellite markers used in this study. The primer sequences were mapped against scaffolds downloaded from ftp://ccb.jhu.edu/pub/data/Redwood/Assembly/v1.0 using the CLC Genomic Workbench v11.0.1 software (Qiagen, Hilden, Germany) with the following parameters: complete match for the last 15 nucleotides at 3’ end, maximum mismatch for 3 nucleotides per annealing site in total, and a maximum of 400 nucleotides between annealing sites.

### Genotyping of the nuclear microsatellite (nSSR) markers

GeneMapper 4.1. (Applied Biosystems) was used for visualization and fragment size calling of the PCR products. *S. sempervirens* is a hexaploid species [38], which complicates microsatellite scoring [33, 39]. In this study, we used the genotyping method of Pfeiffer et al. [40] to identify consistent and reproducible alleles. Following this method, all detected fluorescent peaks representing different PCR amplified fragments were assigned in each sample to one of the 10 rank categories according to the shape and intensity of the peak and the motif of the primer. It was done for all samples representing 30 clones, including 2-3 ramets per clone. To determine which ranking has the highest probability to be correct, input files with different ranking combinations from category 7 to 10 with all 12 primers were built. For example, for the ranking combination 7, all peaks of each sample assigned to the category 7 and above (8, 9 and 10) were included in the analysis, for the ranking combination 8, all peaks of each sample assigned to the category 8 and above (9 and 10) were included into the analysis; etc. The R-package “polysat” [41] was used for each ranking combination data set to calculate the pairwise ‘*bruvo genetic distance’* between individuals [42]. Using the R-package “ape” [43] and the pairwise distance matrix, a Neighbour Joining (NJ) consensus tree was generated based on 1000 bootstraps. Since each marker characteristic differed, 25 different ranking combinations for the markers were tested. The ranking combination with the highest bootstrap values and correct grouping of the ramets representing the same clone was used for all further genotype scoring. The NJ results were confirmed with the function “assignClones” of the R-package “polysat” using the miss-matching threshold of 0.2 [44].

The markers were finally ranked between 7 to 10 (**Error! Reference source not found.**Table). The NJ tree (NJT) based on the final ranking combination of the markers and 1000 bootstraps is presented in S3 Fig.

In addition, for the data sets F and C, scored genotypes were converted also into a presence-absence matrix, and the Nei’s genetic distances ([45] after [46]) were calculated between watersheds F and populations C using the AFLPsurv software with 1000 permutations [47]. The genetic distance matrix was used to generate a NJT with the PHYLIP v369 software [48], respectively. The consensus tree was visualized using the FIGTREE software [49].

### Genotyping of the chloroplast microsatellite (cpSSR) markers and haplotype network

Due to the haploid nature of chloroplasts, genotyping of the cpSSR markers was easier and unambiguous. Based on all three data sets, a haplotype network was built using the Goldstein distance [50] in the program EDENetwork v2.28 [51]. The program calculates a weighted network based on the Goldstein distance between haplotypes with an automatically calculated percolation threshold of 2.67 [52].

### Genetic assignment using the chloroplast microsatellite (cpSSR) markers

The individuals from the German data set G were assigned to the data sets F and C using the GeneClass2 software [53]. To do that eight closely located sampling sites (< 1.0 km) were combined into four. PUC and LPF were combined into LPF (watershed M), MtMa1 and MtMa2 into MtMa (watershed O), HW1 and HW2 into HW (watershed G), and JDF and CCJDF into JDF (watershed J). A quality assessment was based on the self-assignment test following Hintsteiner et al. [16] using the computation criteria of Rannala and Mountain [54], the simulation algorithm of Paetkau et al [55] with 10 000 resampled individuals, and Type 1 error of 0.01 [55, 56]. NJTs were based on Nei’s genetic distance ([45] after [46]) and 1000 bootstraps for the reference data sets C and F and generated using the R-packages “adegenet” [57, 58] and “poppr” [59, 60].

### STRUCTURE analysis

Based either on 12 nSSR or six cpSSR markers input files with the original allele sizes for the data sets F and C were used by the STRUCTURE v2.3.4. program [61–64] to infer the most likely potential number of distinct genetic clusters (*K*) by testing for different number of *K* from 1 to 24 with 20 iterations per run using the MCMC with 10 000 burn in’ and 100 000 final iterations assuming the admixture model. The STRUCTURE HARVESTER [65] and the ClumPPAK [66] programs were used to visualize the STRUCTURE results and to help determining the most likely number of clusters using the Δ*K* approach. The STRUCTURE analyses were repeated also with the “locprior” optional parameter engaged (which is used to assist the clustering by specifying populations a prior), because low population differentiation was expected. Following the STRUCTURE HARVESTER results, the data sets F and C were divided each into two separately analysed groups - 12 northern populations and four southern populations of the San Francisco Bay (south of the Napa Valley) to test for further clustering within these two groups using the same STRUCTURE settings.

## Results

Mapping of the PCR primer nucleotide sequences for the SSR markers against the draft coast redwood nuclear and chloroplast genome assemblies found annealing sites facing each other in a correct configuration allowing amplification for all six chloroplast markers and eight out of 12 nuclear markers (see S4 Table for details). Three different single scaffolds included annealing sites for three markers *ss36782*, *ss73361*, and *ss114481*, respectively, two different scaffolds included annealing sites for each of two markers *RW48* and *ss73307*, respectively, three, four, and six different scaffolds included annealing sites for *ss73978*, *ss91170*, and *ss74800* markers. The chloroplast markers all had only a single annealing site per each marker in the chloroplast genome, but multiple scaffolds also contained annealing sites. The latter could be due to either not complete removal of chloroplast sequences from the nuclear genome assembly or presence of chloroplast sequences in the nuclear genome as a result of misassembling or migration of chloroplast genes to the nuclear genome [67–69].

The NJTs for the data sets F and C had low bootstrap values below 60 % indicating weak differentiation between watersheds and populations based on 12 nSSR markers (Fig 3 and 4), excepts for two watershed pairs P-Q and E-G in the data set F (Fig 3) and populations AET and CCJDF in the data set C (Fig 4). The NJTs based on the cpSSR markers showed higher bootstrap values for most clusters, but they did not reflect their geographic relationship (S4 and S5 Figs).

**Fig 3.**
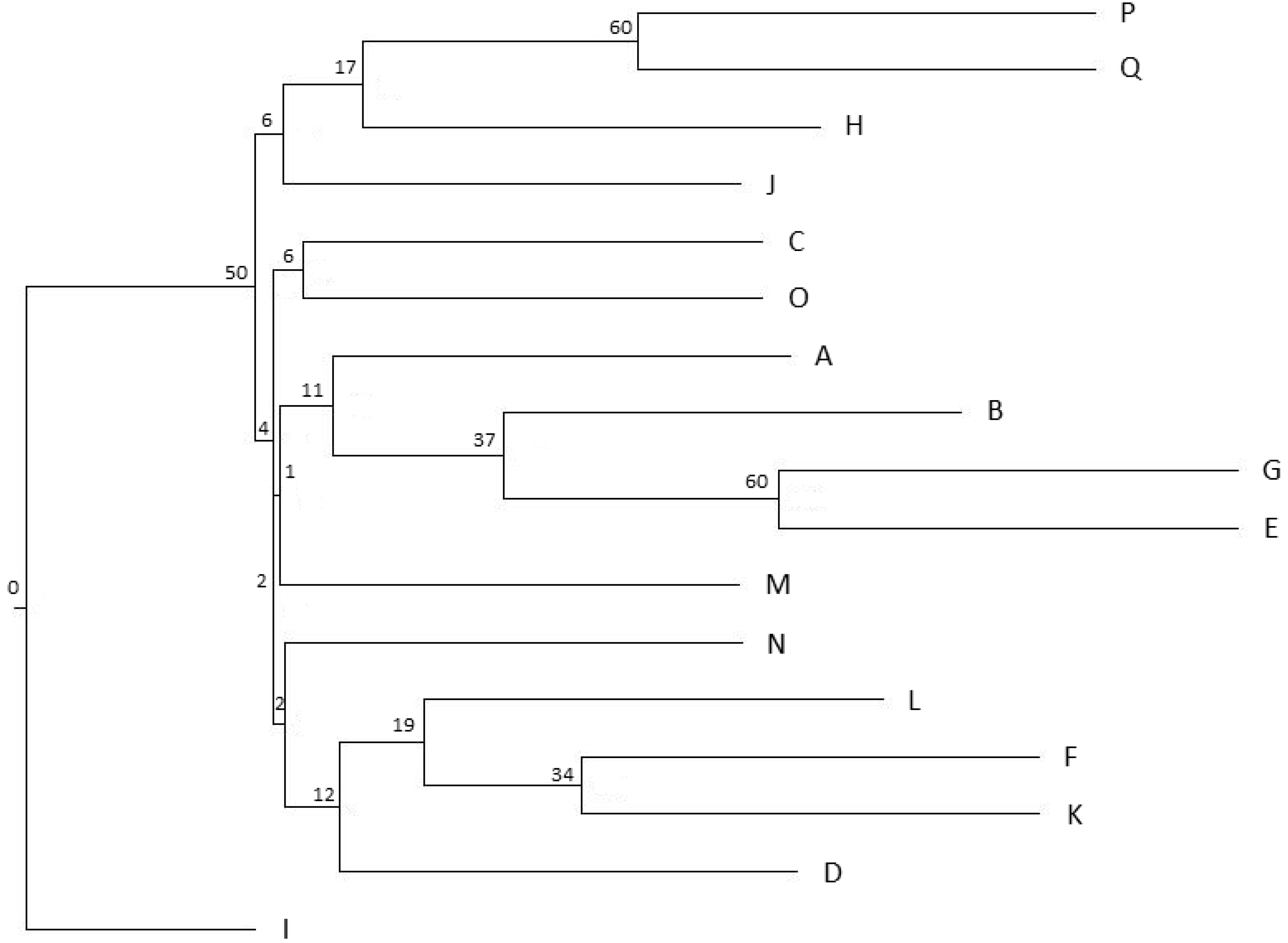
The neighbour-joining tree for the French data set F partitioned into 17 watersheds (A-Q) following Douhovnikoff & Dodd [31] based on the Nei’s genetic distance ([45] after [46]) calculated using 12 nSSR markers. Watersheds from A to Q are distributed from north (Oregon border) to the south of central California (see also Fig 1 and S5 Table). Bootstrap values are presented as percentage.

**Fig 4.**
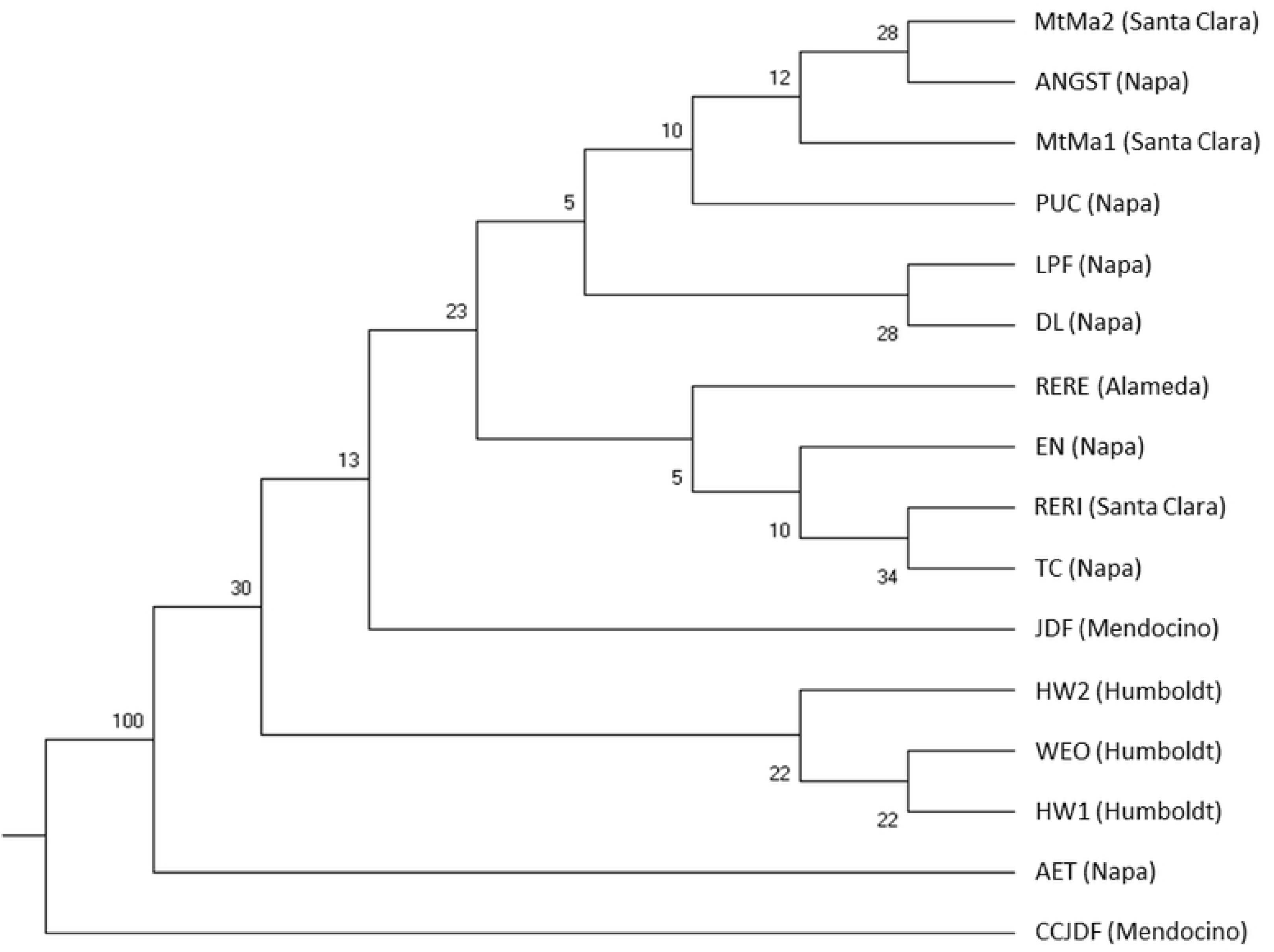
The neighbour-joining tree for populations in the Californian data set C based on the Nei’s genetic distance ([45] after [46]) calculated using 12 nSSR markers. County names of the sampled locations are in brackets. Bootstrap values are presented as percentage.

The STRUCTURE analysis also suggested very low differentiation with maximum two clusters (*K* = 2) for the data sets F and C (Fig. Fig.). Additional STRUCTURE analysis performed separately within the northern and southern subpopulations did not find additional clusters and confirmed that the differentiation observed in the data sets F and C was mainly due to the differences between northern and southern populations. The “locprior” function did not affect much the STRUCTURE results, therefore only results obtained with this function engaged are presented in Fig. Fig. 6, and results obtained with the “locprior” function disengaged can be found in S7 and S8 Figs.

**Fig. 5.**
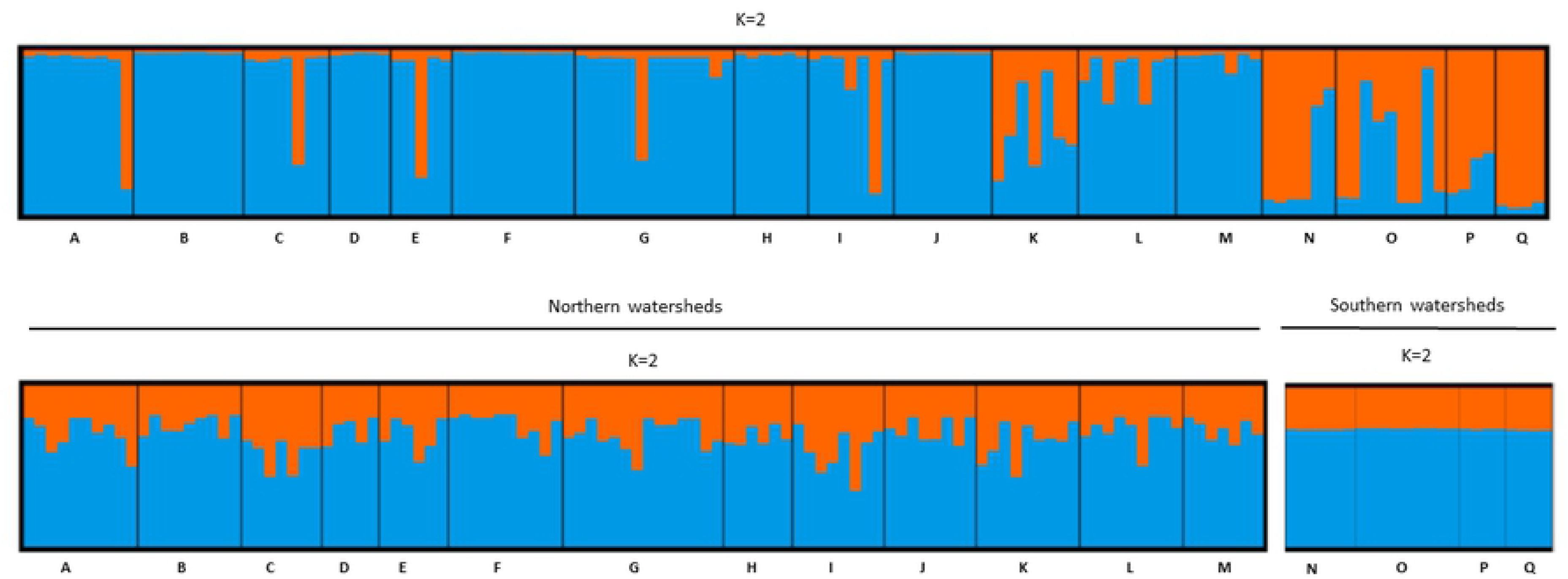
STRUCTURE analysis demonstrating the most likely number of clusters (*K*) using the “locprior” function for 16 californian locations in the data set C based on 6 cpSSR markers, and two separately analysed groups - 12 northern locations (*K* = 3) and four southern locations of San Francisco Bay (*K* = 4).

**Fig. 6.**
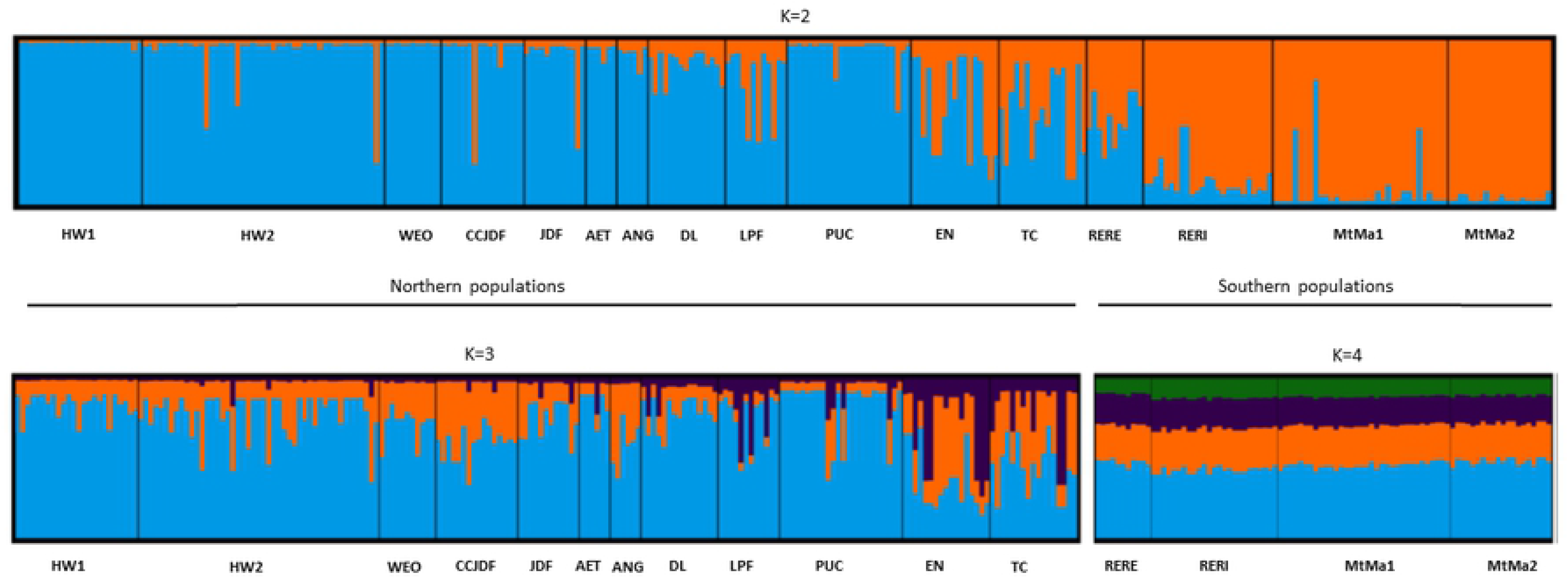
STRUCTURE analysis demonstrating the most likely number of clusters (*K*) using the “locprior” function for the French data set F based on 6 cpSSR markers.

In the data set G, 95 individual trees from the Burgholz stand were identified as likely clones of the SF72 tree from the “Sequoiafarm Kaldenkirchen”. Other individuals from Burgholz and Bad Grund were very similar to the SF76 tree, but with insufficient statistical support to assign them all unambiguously to one clone.

Based on six cpSSR markers, 63 haplotypes in the data set F, 109 in C, and 22 in G were found. All three data sets combined contained 150 different haplotypes (Fig. 7). The haplotype network and its percolated clusters generated by the EDENetwork software indicated no geographic structure in relationships between the haplotypes (Fig.). More than half of all haplotypes were unique - 89 haplotypes occurred only once within the three data sets. The haplotypes GG and KK were the most frequent and were observed with frequency 17 % and 14 %, respectively.

**Fig. 7.**
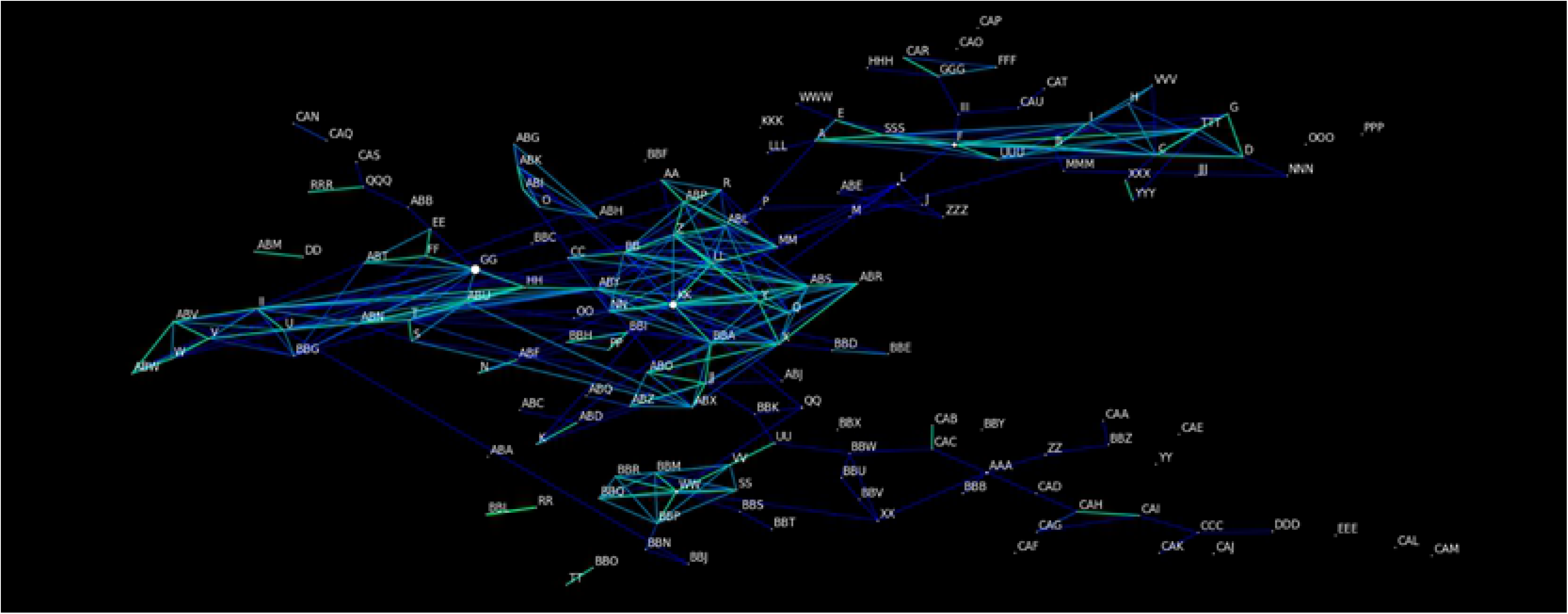
Haplotype network using the percolation threshold of 2.67 based on the cpSSR markers genotyped in three data sets F, C and G. Size of the nodes reflects number of individuals assigned to the respective haplotype. Light green presents closest connections, darker colours indicate decreasing relationship between haplotypes, and can be considered as alternative connections.

Accuracy of the assignments was evaluated by the quality index (QI), which was low for both F (QI= 7 %) and C (QI= 16 %).

Individuals with identical genotype representing the same clone SF71 were presented as a single entry in Table 1.**Table 1. Table 1. Table 1. Table 1.**

**Table 1.** Assigned populations and watersheds and assignment quality scores (%) for two ranks of the German samples to the populations and watersheds in the reference data sets representing samples from California (C) and the “Kuser Provenance test” in St. Fargeau, central France (F), respectively.

| Sample | Reference C |  |  |  | Reference F |  |  |  |
| --- | --- | --- | --- | --- | --- | --- | --- | --- |
|  | Rank 1 |  | Rank 2 |  | Rank 1 |  | Rank 2 |  |
|  | Assigned populations and watersheds (county) | Score, % | Assigned populations and watersheds (county) | Score, % | Assigned watersheds (county) | Score, % | Assigned watersheds (county) | Score, % |
| GOE_P | RERI, O (Santa Clara) | 97 | RERE, N (Santa Clara) | 2 | Q (Monterey) | 55 | N (Marine) | 14 |
| B13 | JDF, J (Mendocino) | 86 | AET, M (Napa) | 6 | G (Humboldt) | 33 | C (Del Norte) | 26 |
| B42 | JDF, J (Mendocino) | 86 | AET, M (Napa) | 6 | G (Humboldt) | 33 | C (Del Norte) | 26 |
| B51 | JDF, J (Mendocino) | 86 | AET, M (Napa) | 6 | G (Humboldt) | 33 | C (Del Norte) | 26 |
| B78 | JDF, J (Mendocino) | 86 | AET, M (Napa) | 6 | G (Humboldt) | 33 | C (Del Norte) | 26 |
| SF85 | JDF, J (Mendocino) | 86 | AET, M (Napa) | 6 | G (Humboldt) | 33 | C (Humboldt) | 26 |
| BA10067 | MtMa, O (Santa Clara) | 85 | LPF, M (Napa) | 6 | O (Santa Cruz) | 60 | N (Marine) | 15 |
| GOE_B | MtMa, O (Santa Clara) | 65 | RERI, O (Santa Clara) | 15 | P (Monterey) | 69 | K (Mendocino) | 18 |
| SF65 | WEO, G (Humboldt) | 59 | HW, G (Humboldt) | 14 | E (Humboldt) | 30 | I (Mendocino) | 27 |
| SF80 | AET, M (Napa) | 58 | JDF, J (Mendocino) | 16 | J (Mendocino) | 39 | F (Humboldt) | 32 |
| SF75 | JDF, J (Mendocino) | 58 | TC, M (Napa) | 30 | J (Mendocino) | 25 | G (Humboldt) | 22 |
| SF67 | JDF, J (Mendocino) | 52 | AET, M (Napa) | 20 | D (Humboldt) | 45 | E (Humboldt) | 14 |
| SF70 | HW, G (Humboldt) | 50 | EN, M (Napa) | 35 | F (Humboldt) | 74 | B (Del Norte) | 5 |
| SF74 | EN, M (Napa) | 48 | HW, G (Humboldt) | 19 | G (Humboldt) | 33 | B (Del Norte) | 8 |
| SF81 | RERE, N (Santa Clara) | 47 | EN, M (Napa) | 18 | C (Humboldt) | 38 | A (Del Norte) | 28 |
|  | Rank 1 |  | Rank 2 |  | Rank 1 |  | Rank 2 |  |
|  | Assigned populations and watersheds (county) | Score, % | Assigned populations and watersheds (county) | Score, % | Assigned watersheds (county) | Score, % | Assigned watersheds (county) | Score, % |
| SF73 | JDF, J (Mendocino) | 44 | HW, G (Humboldt) | 13 | B (Del Norte) | 14 | J (Mendocino) | 12 |
| B4 | AET, M (Napa) | 43 | HW, G (Humboldt) | 15 | H (Mendocino) | 19 | M (Napa) | 16 |
| CH1 | AET, M (Napa) | 43 | HW, G (Humboldt) | 15 | H (Mendocino) | 19 | M (Napa) | 16 |
| SF3 | AET, M (Napa) | 43 | HW, G (Humboldt) | 15 | H (Mendocino) | 19 | M (Napa) | 16 |
| SF63 | AET, M (Napa) | 43 | HW, G (Humboldt) | 15 | H (Mendocino) | 19 | M (Napa) | 16 |
| SF64 | AET, M (Napa) | 43 | HW, G (Humboldt) | 15 | H (Mendocino) | 19 | M (Napa) | 16 |
| SF76 | AET, M (Napa) | 43 | HW, G (Humboldt) | 15 | H (Mendocino) | 19 | M (Napa) | 16 |
| SF79 | AET, M (Napa) | 43 | HW, G (Humboldt) | 15 | H (Mendocino) | 19 | M (Napa) | 16 |
| SF82 | AET, M (Napa) | 43 | HW, G (Humboldt) | 15 | H (Mendocino) | 19 | M (Napa) | 16 |
| SF86 | AET, M (Napa) | 43 | HW, G (Humboldt) | 15 | H (Mendocino) | 19 | M (Napa) | 16 |
| SF87 | AET, M (Napa) | 43 | HW, G (Humboldt) | 15 | H (Mendocino) | 19 | M (Napa) | 16 |
| SF93 | AET, M (Napa) | 43 | HW, G (Humboldt) | 15 | H (Mendocino) | 19 | M (Napa) | 16.297 |
| BG120 | JDF, J (Mendocino) | 38 | HW, G (Humboldt) | 33 | F (Humboldt) | 29 | B (Del Norte) | 17.869 |
| SF84 | JDF, J (Mendocino) | 35 | DL (Napa) | 24 | J (Mendocino) | 30 | M (Napa) | 23 |
| SF91 | DL (Napa) | 33 | RERE, N (Santa Clara) | 32 | B (Del Norte) | 23 | N (Marine) | 18 |
| SF69 | JDF, J (Mendocino) | 31 | AET, M (Napa) | 31 | F (Humboldt) | 41 | H (Mendocino) | 15 |
| SF88 | JDF, J (Mendocino) | 31 | WEO, G (Humboldt) | 19 | L (Sonoma) | 25 | A (Del Norte) | 18 |
|  | Rank 1 |  | Rank 2 |  | Rank 1 |  | Rank 2 |  |
|  | Assigned populations and watersheds (county) | Score, % | Assigned populations and watersheds (county) | Score, % | Assigned watersheds (county) | Score, % | Assigned watersheds (county) | Score, % |
| SF90 | JDF, J (Mendocino) | 31 | WEO, G (Humboldt) | 19 | L (Sonoma) | 25 | A (Del Norte) | 18 |
| BA030549 | RERE, N (Santa Clara) | 28 | DL (Napa) | 24 | A (Del Norte) | 31 | D (Humboldt) | 9 |
| B30 | MtMa, O (Santa Clara) | 25 | RERI, O (Santa Clara) | 25 | O (Santa Cruz) | 24 | L (Sonoma) | 17 |
| B89 | MtMa, O (Santa Clara) | 25 | RERI, O (Santa Clara) | 25 | O (Santa Cruz) | 24 | L (Sonoma) | 17 |
| B102 | LPF, M (Napa) | 21 | HW, G (Humboldt) | 17 | B (Del Norte) | 14 | L (Sonoma) | 12 |
| BG121 | JDF, J (Mendocino) | 21 | HW, G (Humboldt) | 17 | B (Del Norte) | 14 | L (Sonoma) | 12 |
| BG122 | JDF, J (Mendocino) | 21 | HW, G (Humboldt) | 17 | B (Del Norte) | 14 | L (Sonoma) | 12 |
| BG123 | JDF, J (Mendocino) | 21 | HW, G (Humboldt) | 17 | B (Del Norte) | 14 | L (Sonoma) | 12 |
| BG124 | JDF, J (Mendocino) | 21 | HW, G (Humboldt) | 17 | B (Del Norte) | 14 | L (Sonoma) | 12 |
| BG125 | JDF, J (Mendocino) | 21 | HW, G (Humboldt) | 17 | B (Del Norte) | 14 | L (Sonoma) | 12 |
| BG126 | JDF, J (Mendocino) | 21 | HW, G (Humboldt) | 17 | B (Del Norte) | 14 | L (Sonoma) | 12 |
| BG127 | JDF, J (Mendocino) | 21 | HW, G (Humboldt) | 17 | B (Del Norte) | 14 | L (Sonoma) | 12 |
| BG128 | JDF, J (Mendocino) | 21 | HW, G (Humboldt) | 17 | B (Del Norte) | 14 | L (Sonoma) | 12 |
| GOE_G | JDF, J (Mendocino) | 21 | HW, G (Humboldt) | 17 | B (Del Norte) | 14 | L (Sonoma) | 12 |
| GOE_K | JDF, J (Mendocino) | 21 | HW, G (Humboldt) | 17 | B (Del Norte) | 14 | L (Sonoma) | 12 |
| SF62 | JDF, J (Mendocino) | 21 | HW, G (Humboldt) | 17 | B (Del Norte) | 14 | L (Sonoma) | 12 |
| SF66 | JDF, J (Mendocino) | 21 | HW, G (Humboldt) | 17 | B (Del Norte) | 14 | L (Sonoma) | 12 |
|  | Rank 1 |  | Rank 2 |  | Rank 1 |  | Rank 2 |  |
|  | Assigned populations and watersheds (county) | Score, % | Assigned populations and watersheds (county) | Score, % | Assigned watersheds (county) | Score, % | Assigned watersheds (county) | Score, % |
| SF83 | JDF, J (Mendocino) | 21 | HW, G (Humboldt) | 17 | B (Del Norte) | 14 | L (Sonoma) | 12 |
| B11 | RERI, O (Santa Clara) | 19 | AET, M (Napa) | 15 | B (Del Norte) | 13 | L (Sonoma) | 11 |
| B35 | RERI, O (Santa Clara) | 19 | AET, M (Napa) | 15 | B (Del Norte) | 13 | L (Sonoma) | 11 |
| B89a | RERI, O (Santa Clara) | 19 | AET, M (Napa) | 15 | B (Del Norte) | 13 | L (Sonoma) | 11 |
| SF72 | RERI, O (Santa Clara) | 19 | AET, M (Napa) | 15 | B (Del Norte) | 13 | L (Sonoma) | 11 |
| SF77 | RERI, O (Santa Clara) | 19 | AET, M (Napa) | 15 | B (Del Norte) | 13 | L (Sonoma) | 11 |
| SF68 | JDF, J (Mendocino) | 40 | AET, M (Napa) | 17 | F (Humboldt) | 44 | H (Mendocino) | 8 |
| SF71 | RERI, O (Santa Clara) | 21 | TC, M (Napa) | 20 | O (Santa Cruz) | 47 | I (Mendocino) | 13 |

GOE - Forest Botanical Garden of the University of Göttingen, B - Burgholz, BA - Ecological-Botanical Garden of the University of Bayreuth, BG - Bad Grund, CH - Botanical Garden of Chemnitz, SF - Sequoiafarm Kaldenkirchen. The blue and yellow colors highlight the two clusters suggested by STRUCTURE, north and south of the San Francisco Bay, respectively. In brackets are the names of the counties for the respective locations (C) and watersheds (F).

Most of the assignments of German trees to the southern or northern populations divided by the San Francisco Bay in the reference data set C with scores above 48 % were in agreement with the assignments in the reference data set F (**Error! Reference source not found.**). While some samples with scores below 48 % were assigned to geographically very different watersheds. For example, B11 and SF77 were assigned to RERI south of the San Francisco Bay (watershed O) based on the reference data set C, but to Del Norte close to the Oregon border (watershed B) based on reference F (Table 1, **Error! Reference source not found.**). The individuals from the Sequoiafarm, which were half- or full siblings according to the owner, were assigned to various watersheds (F) and locations (C).

## Discussion

The low population differentiation within the natural distribution range of *S. sempervirens* based on the supposedly selectively neutral nuclear and chloroplast markers and both reference data sets F and C, respectively, confirmed results of previous studies [24,25,31]. The NJT for the data set F based on 12 nSSR markers in our study was in consensus with the NJT calculated for the same 12 watersheds (A-Q) by Douhovnikoff and Dodd [31] using clones from the Russell Reserve. To facilitate comparison between these two phylogenetic trees, the watershed I (Mendocino County) was also used as a root of the NJT. The discrepancies between the two trees could be explained by very low bootstrap support for most clusters in both trees and by using the pairwise *F*_ST_ values for clustering in Douhovnikoff and Dodd [31] instead of the Nei’s genetic distance used in our study. Moreover, the calculations in Douhovnikoff and Doddwere based on less than half of nSSR markers (6 vs. 12), and only two of these markers were used in both studies.

The STRUCTURE analyses based on 12 nSSR and six cpSSR markers confirmed the NJT results of low differentiation, but analyses based on the cpSSR markers were able to identify the San Francisco Bay as a border between two main clusters of populations. However, the obtained results also had only low statistical support (S7 and S8 Figs). The STRUCTURE analyses within each of the two areas, north and south of the Bay, did not reveal any additional clusters, neither in the dataset F nor C. Our data confirmed the San Francisco Bay as a border suggested already earlier by Brinegar [30] based on a single chloroplast marker. It is in consensus also with one of two borders identified by Sawyer et al. [70] based on the soil conditions and water availability provided by precipitation and fog.

The lack of strong population structure and low genetic differentiation can explain the inconsistent assignment of German trees to populations in both reference data sets C and F and the low QI for them. For reliable assignment a stronger differentiation between populations in a reference data set and sufficient sample size of each reference population are needed [15,71,72]. Sample sizes were possibly insufficient for some populations in the reference C and F ranging from 4 to 47 individuals per population or watershed. However, considering the results of the STRUCTURE analyses that suggested northern and southern clusters, all but five German trees were assigned correctly to these clusters (**Error! Reference source not found.**). The potential errors in the records on the origin of trees in the reference populations need also to be taken into account when considering the reliability of the assignment of individuals to an origin, because individuals with wrongly identified origin within the reference population can decrease the assignment quality [56]. The possibility of trees with wrongly identified origin and non-local genotypes being included in the two reference sets was quite high because coast redwood is a heavily used timber species with a long tradition of planting, replanting, and plant material transfer [73]. Trees from areas north of San Francisco Bay might have been planted in the south and vice versa. This is really hard to detect and could only be verified in a comprehensive and detailed population genetic study of natural coast redwood populations in California and Oregon.

The reliable clone identification in the data set G (S6 Fig) confirmed results of previous studies based on allozyme and AFLP markers [24, 74].

General difficulties to accurately genotype microsatellite markers in polyploid organisms excludes analyses based on their allele frequencies [33,75–78]. It concerns also coast redwood, but genotyping problems could be even more aggravated due to a high probability of somatic mutations in basal sprouting shoots in these extremely long living trees, which can result in different genotypes of different tissues and clones originated from the same tree [33]. The assumption of Hardy-Weinberg equilibrium is also tricky due to common clonal growth in coast redwood populations [66], where trees within 40 m distances can belong to the same clone [74]. The correct estimation of the null allele frequencies and allelic dosage are two major difficulties associated with genotyping polyploidy organisms using microsatellite markers [79]. In our study the risk of null alleles should be reduced since null alleles are in general less frequent in EST-SSRs due to their location in more conserved sequences [79], although they are still possible and could affect the assignment results in this study. The Bruvo distance used as the genetic distance measure in this study does not require same allelic dosages between individuals [41, 42]. The fragment scoring following Pfeiffer et al. [40] and confirmation of allele scoring using multiple ramets of the same clones reduced the scoring errors due to the stutter effects in this study. Narayan et al. [33] considered only those alleles for the multilocus lineages (MLL) that were consistent in at least two different tissue types of the same tree, and they found a low error rate in allelic dosage in coast redwood based on the Bruvo genetic distance. The possible number of genotypes in highly polyploid species, such as coast redwood, can be very high [40, 72], hence, only few microsatellite primers can be sufficient for clonal identification and population structure analysis.

The number of chloroplast haplotypes found in this study was exceptionally high (150) based on six cpSSR markers genotyped in 579 samples in all data sets F, C and G combined. It confirmed higher variation of cpSSR than nSSR markers usually observed in conifers [80], although the mutation rate of chloroplast microsatellite loci is lower [81]. However, cluster analysis did not reveal geographic differentiation other than two northern and southern groups of populations. Similar results were observed in another conifer species, *Abies nordmanniana*, which also had a similar high number of haplotypes, 111, genotyped in 361 individuals, although the sampling range included a several times larger area than for coast redwood [82]. Meanwhile, it should be noted that haplotype differentiation usually reflected the geographic origin of populations in other conifer species [83–85]. However, the differentiation between populations based on the cpSSR markers is usually less than the one based on nSSRs, which can be explained by the paternal inheritance of chloroplasts in *S. sempervirens* [32] and the long distance gene flow via pollen. Ribeiro et al. [86] found similar results when compared population differentiation based on AFLP markers with the one based on the paternally inherited cpSSR markers in the wind pollinated conifer *Pinus pinaster*. In addition, Petit et al. [87] showed that for various conifer species genetic differentiation based on bi-parentally inherited markers correlated with differentiation based on paternally inherited markers due to the similar gene flow vectors, but the latter one was in general lower [88].

## Conclusions and future directions

Coast redwood forest used to have continuous distribution along the pacific coast in California before intensive logging started at the beginning of the nineteenth century [89]. Therefore, current coast redwood populations are considered as remnant and fragmented populations. However, being long living clonal trees with high somatic mutation rate coast redwood maintained its high genetic diversity despite multiple bottlenecks [8, 32].

The combination of low sexual reproduction and local adaptation that could be insufficient to meet the predicted future climate change will increase the pressure on coast redwood [8]. California has become drier in the last 2000 years [9], and the very important fog has been declining in its frequency during the last century [36, 90]. Considering these threats O’Hara et al. [8] emphasized the necessity to find drought tolerant genotypes, especially for southern populations. Genetic and physiological mechanisms behind drought resistance are similar to those that are behind frost tolerance [91]. Geographic variation in drought tolerance in tree species often overlaps with its variation in frost tolerance because the physiological mechanisms of drought tolerance and frost tolerance are similar [92]. One of the mechanisms of frost resistance is related to preventing crystal water formation via osmotic regulation of the cells, which is also important in drought resistance [93]. The identification of water stress resistant genotypes would benefit both Californian and German forestry. Tolerant genotypes would not only provide Germany with a valuable timber species considering climate change, but also presents suitable resources for *ex-situ* conservation programs for coast redwood, which was already suggested for the sister species *Sequoiadendron giganteum* [2].

Further studies of the genetic structure of coast redwood populations and clones are needed based on functional markers that are potentially under selection, such as non-synonymous SNPs or SNPs in regulatory genes. The results obtained in these studies would provide important data for sustainable timber production and improved conservation management, *in situ* and *ex situ*, for coast redwood, especially in the context of predicted climate change.

## Supporting information

**S1 Table. Watershed information.** Latitude and number of samples representing 17 watersheds in the French (F) data set (St. Fargeau) and the Russell reserve according to Douhovnikoff and Dodd [31].

**S2 Table. Geographic and climatic data for the locations sampled in California (data set C).**

**S3 Table. Data for 18 microsatellite (SSR) markers used in the study.**

**S4 Table. Verification of the 18 PCR primer pairs of the 18 microsatellite (SSR) markers by mapping their primer nucleotide sequences to the coast redwood genome and chloroplast assemblies.**

**S5 Table. ID and geographic data for the French (F) set of the “Kuser’s” samples.**

**S1 Fig. Map of the original “Kuser’s” samples (data set F).** Original locations of the “Kuser’s” samples placed on the map with the mean monthly temperature pattern for the time period 1979-2013 indicating colder and warmer temperatures by darker and lighter shades of grey (http://chelsa-climate.org).

**S2 Fig. Map of the German locations (data set G).** Locations of the samples in the German data set (G) of samples: Sequoiafarm Kaldenkirchen, Arboretum Burgholz, Weltwald Bad Grund, and the Botanic Gardens in Göttingen, Bayreuth and Chemnitz placed on the map with the mean monthly temperature pattern for the time period 1979-2013 indicating colder and warmer temperatures by darker and lighter shades of grey (http://chelsa-climate.org).

**S3 Fig. Neighbour-joining tree of 30 clones collected in the Allerweltsgrün nusery (Köln).** It is based on 12 nSSR markers with the final ranking combination obtained according to Pfeiffer et al. [40] and the ‘bruvo genetic distance’ [42]. Numbers indicate bootstrap values (not percentage).

**S4 Fig. Neighbour-joining tree of the “Kuser’s” samples in the French (F) data set (St. Fargeau) grouped into 17 watersheds (A-Q).** It is based on six cpSSR markers and Nei’s genetic distance ([45] after [46]) with 1000 bootstraps. Numbers indicate bootstrap values in terms of percentage.

**S5 Fig. Neighbour-joining tree of the 16 Californian reference populations represented by samples collected in 2017 (data set C).** It is based on six cpSSR markers and Nei’s genetic distance ([45] after [46]) with 1000 bootstraps. Numbers indicate bootstrap values in terms of percentage. County names of sampling origin are in brackets.

**S6 Fig. Neighbour-joining tree of the 143 individual trees collected in Germany (data set G).** It is based on 12 nSSR markers with the final ranking combination obtained according to Pfeiffer et al. [40] and the ‘bruvo genetic distance’ [42] with 1000 bootstraps. Numbers indicate bootstrap values (not percentage).

**S7 Fig. STRUCTURE results for the reference data sets C and F based on 12 nSSR markers.** Plots for data sets C and F are presented for the most likely number of clusters (*K*) inferred without (a) and with (b) the “locprior” function (*K* = 6 and *K* = 2 for C and *K* = 2 for F, respectively). L(*K*) and Δ*K* statistics generated by the ClumPAK software are also presented.

**S8 Fig. STRUCTURE statistics for the reference data sets C and F based on 6 cpSSR markers.** Presented are the most likely number of clusters (*K*) and results of STRUCTURE runs for *K* from 1 to 20 or 25 with 20 iterations each for the complete data set C (a), northern subset C (b), southern subset C (c), complete data set F (d), northern subset F (e), and southern subset F (f).

